# Enhanced virus translation enables miR-122-independent Hepatitis C Virus propagation

**DOI:** 10.1101/2021.06.08.447644

**Authors:** Mamata Panigrahi, Michael A Palmer, Joyce A Wilson

**Affiliations:** Department of Biochemistry, Microbiology and Immunology, College of Medicine, University of Saskatchewan, Saskatoon, SK, Canada

**Keywords:** HCV, miR-122-independent replication, viral translation, 5’ UTR, genome stability

## Abstract

The 5’UTR of the Hepatitis C Virus genome forms RNA structures that regulate virus replication and translation. The region contains a viral internal ribosomal entry site and a 5’ terminal region. Binding of the liver specific miRNA, miR-122, to two conserved binding sites in the 5’ terminal region regulates viral replication, translation, and genome stability, and is essential for efficient virus replication, but its precise mechanism of its action is still under debate. A current hypothesis is that miR-122 binding stimulates viral translation by facilitating the viral 5’ UTR to form the translationally active HCV IRES RNA structure. While miR-122 is essential for detectable virus replication in cell culture, several viral variants with 5’ UTR mutations exhibit low level replication in the absence of miR-122. We show that HCV mutants capable of replicating independently of miR-122 also replicate independently of other microRNAs generated by the canonical miRNA synthesis pathway. Further, we also show that the mutant genomes display an enhanced translation phenotype that correlates with their ability to replicate independently of miR-122. Finally, we provide evidence that translation regulation is the major role for miR-122, and show that miR-122-independent HCV replication can be rescued to miR-122-dependent levels by the combined impacts of 5’ UTR mutations that stimulate translation, and by stabilizing the viral genome by knockdown of host exonucleases and phosphatases that degrade the genome. Thus, we provide a model suggesting that translation stimulation and genome stabilization are the primary roles for miR-122 in the virus life cycle.

**IMPORTANCE:** The unusual role of miR-122 in promoting HCV propagation is incompletely understood but is essential for an HCV infection. To better understand its role, we have analyzed HCV mutants capable of replicating independently of miR-122. Our data show that the ability of viruses to replicate independently of miR-122 correlates with enhanced virus translation, but that genome stabilization is required to restore efficient HCV replication. This suggests that viruses must gain both abilities to escape the need for miR-122 and impacts the possibility that HCV can evolve to replicate outside of the liver.

## INTRODUCTION

Hepatitis C virus (HCV) chronically infects almost 71 million people worldwide (1) and can lead to liver cirrhosis and hepatocellular carcinoma (2, 3). HCV has a positive-strand RNA genome 9.6 kb in length that contains a polyprotein coding region flanked by the 5’ and 3’ untranslated regions (UTRs). The 5’ UTR of the viral genome is highly structured and forms several stem-loops (SL), from SLI-SLIV (4–6) that modulate viral replication and translation (7). The 5’ terminal 1-42 nucleotides comprise stem-loop 1 (SLI) and two binding sites (S1 and S2) for a host liver-specific microRNA, miR-122, whose annealing is required for efficient viral propagation (8–12). Nucleotides 40 to 372 of the viral 5’ UTR including SLII to SLIV comprise the HCV internal ribosomal entry site (IRES) that directs cap-independent virus translation initiation (13–15). The complement of the 5’ terminal region, the 3’ UTR of the negative strand, also contains RNA stem-loops that are important for viral replication (6, 16). Thus, the extreme 5’ end of the viral genome is a multifunctional regulatory sequence that harbors structures required for both viral replication and translation.

MicroRNAs are short, non-coding RNAs of 20-24 nucleotides in length, that normally downregulate cellular gene expression and facilitate RNA decay (17–19). During miRNA suppression, an endogenously expressed miRNA, in association with one of the host Argonaute proteins (Ago 1-4) forms the core element of the RNA induced silencing complex (RISC) and targets mRNAs for translation silencing and degradation through sequence complementarity within the 3’ UTR (20). However, in an unusual role for a miRNA, annealing of miR-122 to the 5’ UTR of the HCV genome promotes viral propagation and RNA accumulation (10, 11). Proposed functions for miR-122 include stimulation of virus translation (21–23), protecting the viral genome from cellular exonucleases (XRN1 and XRN2), and pyrophosphatases DOM3Z and DUSP11 (24–28), and directly promoting viral genomic RNA replication (29). However, the relative impact of each function on overall viral propagation remains unknown (30).

An emerging hypothesis suggests a role for miR-122 as an RNA chaperone that modulates the structure of the 5’ UTR and the activity of the HCV IRES (11, 12, 31, 32). RNA structure predictions suggest that nucleotides 1-42 preceding viral IRES may modify the thermodynamics of IRES structure formation and that annealing of miR-122 may shift the thermodynamics to favor the active IRES structure formation (**Fig. 1**). While this hypothesis remains to be confirmed using biophysical or biochemical methods, indirect support stems from the analysis of HCV mutants capable of miR-122-independent replication (11, 12). The first model of miR-122-independent replication was a bicistronic subgenomic RNA that contains an EMCV IRES and suggested that altered translation regulation alleviates the need for miR-122 for viral propagation (33). More full-length HCV genomes having mutations/substitution to the 5’ terminal region have been found that exhibit low levels of miR-122-independent replication, but are still dependent on miR-122 for wild-type levels of replication (11, 34–38). The mechanisms of miR-122-independent replication of these HCV variants are still unknown. Prediction software suggests that the 5’ UTR RNA of the mutants has a greater propensity to form the active HCV IRES even in the absence of miR-122 (11, 12) (**Fig. 1**), but a recent report has also highlighted the possibility of binding of other microRNAs to the 5’ UTR of the viral genome (39).

**FIG 1:**
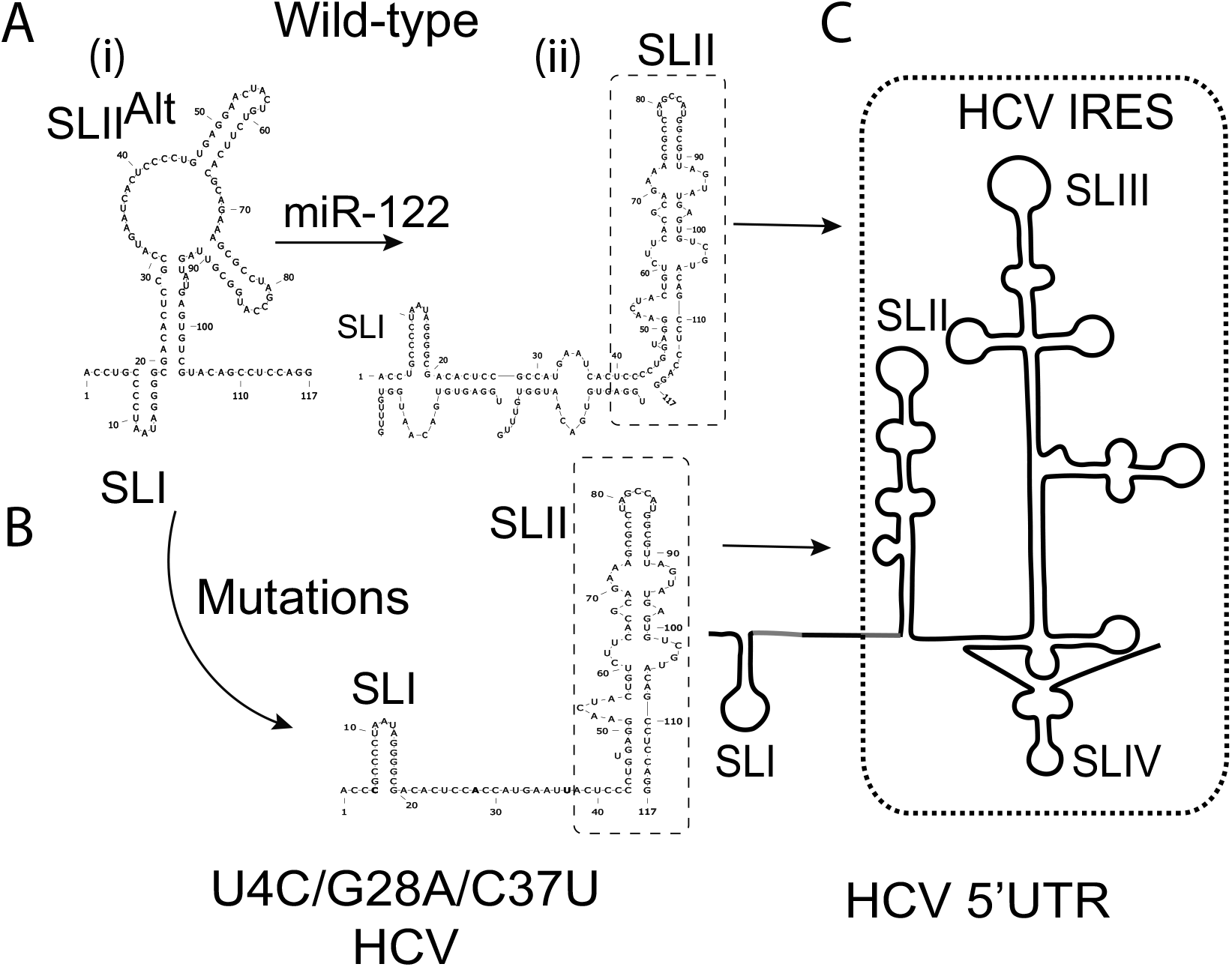
Hypothetical model for miR-122 and 5’ UTR mutation induced RNA structure modifications leading to IRES activation (A) Structure prediction of J6/JFH-1 p7Rluc2a HCV (i) without and (ii) with miR-122. Without miR-122, the first 117 nucleotides of HCV 5’UTR are predicted to form a non-canonical structure with SLI and an altered SLII (SLII^Alt),^ whereas miR-122 annealing to the viral 5’ UTR is predicted to form the canonical SLII structure. (B) The 5’ UTR of a mutant HCV (U4C/G28A/C37U) capable of miR-122-independent replication is also predicted to form the canonical SLII structure in the absence of miR-122. (C) The structure of the active HCV IRES including SLII.

Herein, we have used mutant HCV variants capable of replicating in the absence of miR-122 to study the mechanism of miR-122-independent replication. Our results suggest that HCV mutants capable of replicating independently of miR-122 can also replicate independently of other miRNAs generated by Drosha and the canonical miRNA biogenesis pathways. We also show that mutants capable of miR-122-independent replication display enhanced translation activity compared to the wild-type HCV in the absence of miR-122 and that the translation efficiency correlates with miR-122-independent replication efficiency. Finally, we have characterized the relative contributions of translation and genome stability on miR-122-independent HCV replication and found that translation stimulation is essential for HCV propagation, and enhanced translation and genome stabilization can restore wild-type levels of HCV replication in the absence of miR-122. Thus, our data support that miR-122 has important roles in both viral translation regulation and genome stabilization.

## RESULTS

### HCV mutants capable of miR-122-independent replication can replicate in Drosha knockout cells

Although miR-122 is essential for HCV propagation, we and others have previously reported full-length HCV variants with point mutations in the 5’ UTR of the genome are capable of replicating in the absence of miR-122 (11, 34, 36, 37). However, it is still unclear whether these mutants are capable of usurping other microRNAs in the cell for their propagation. A recent report suggests that HCV replication can be supported by other miRNAs in a miR-122-like (miR-504-3p, miR-574-5p, and miR-1236-5p) and non-miR-122-like manner (miR-25-5p and miR-4730) (39). Thus, we tested for the impact of other miRNAs on the replication of HCV mutants previously reported as capable of replicating independently of miR-122 by assessing their replication in Huh 7.5 Drosha knockout (KO) cells (**Fig. 2A**). This cell line is devoid of Drosha, and thus also devoid of cellular miRNAs generated by the canonical Drosha-dependent biogenesis pathway. Huh7.5 Drosha KO cells were electroporated with wild-type and mutant HCV RNAs, with and without the addition of miR-122. Cell extracts were harvested at 2hr, 24hr, 48hr, and 72hr post electroporation, and luciferase expression was measured as a proxy for viral RNA accumulation. Our results showed that most of the mutants previously reported to replicate independently of miR-122 also replicated in Huh 7.5 Drosha KO cells and thus independently of Drosha-dependent microRNAs, including all the ones identified by Ono *et al* (39)(**Fig. 2B)**. The replication of each mutant in Huh 7.5 Drosha KO cells was comparable to their replication in Huh 7.5 miR-122 knockout (KO) cells. The mutants that grew least efficiently in Huh 7.5 miR-122 KO cells, C26del and C30U/A34G, did not show detectable replication in Huh 7.5 Drosha KO cells, and mutants found previously to display the most efficient miR-122-independent replication, U25C and the triple mutant U4C/G28A/C37U, replicated the most efficiently in Huh 7.5 Drosha KO cells. The G28 series of mutants, excluding G28U, replicated significantly better than the wild-type HCV in the absence of miR-122 in Huh 7.5 Drosha KO cells, similar to their replication in Huh 7.5 miR-122 KO cells. G28U is capable of replicating independently of miR-122 in Huh 7.5 miR-122 KO cells but is less efficient than the other G28 mutants, and the levels were not statistically significantly higher than the wild-type virus.

**FIG 2:**
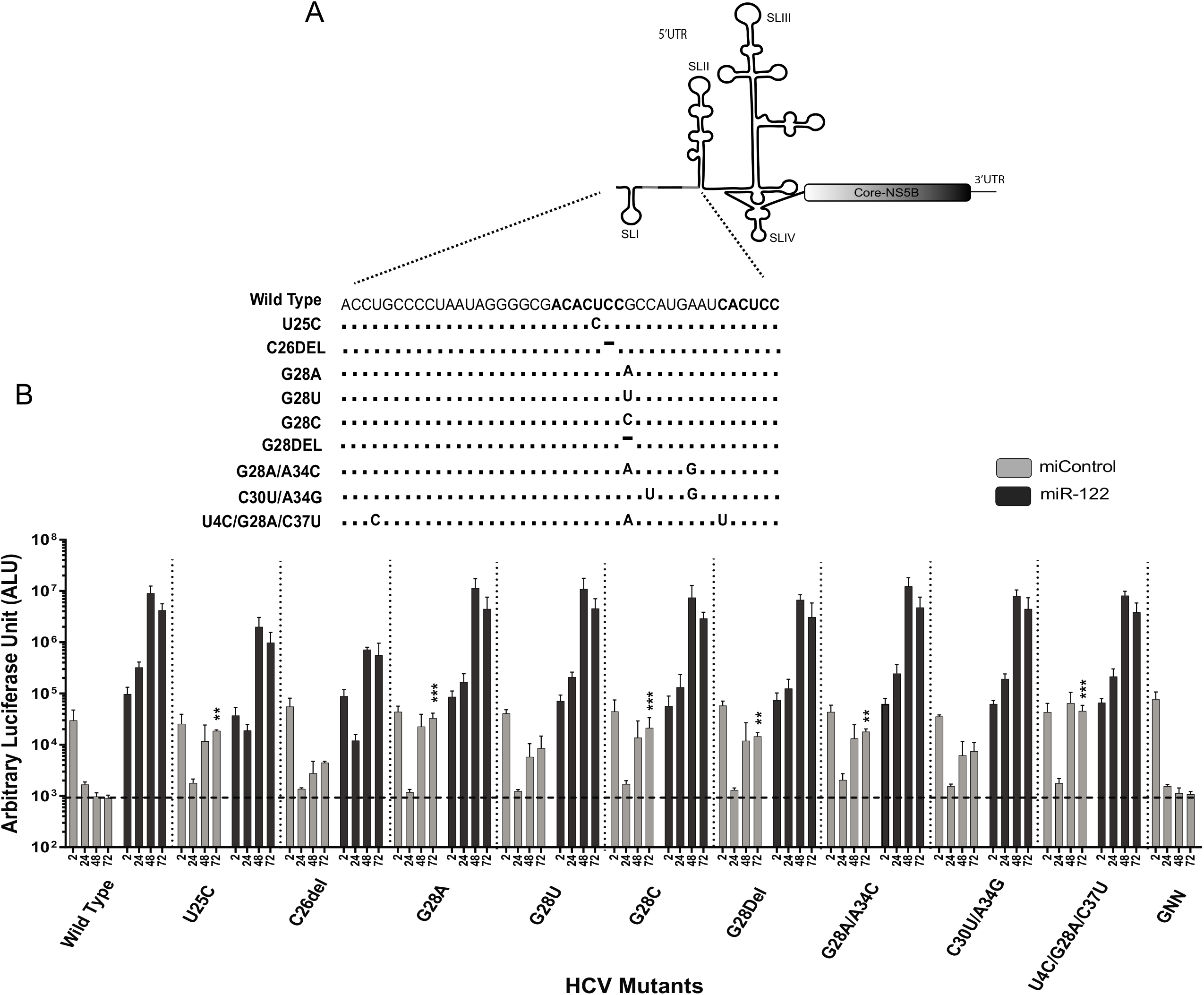
Replication of HCV mutants in Huh 7.5 Drosha knockout cells: (A) Reported mutations on the extreme 5’ UTR of the viral genome that promote virus replication independently of miR-122. The number indicates the position of the nucleotide with the mutation. (B) Transient replication of HCV mutants in Huh 7.5 Drosh knockout cells electroporated with J6/JFH-1 p7Rluc2a HCV RNA with a control miRNA (grey bars) or miR-122 (black bars). Cells were harvested at 2h, 24h, 48h, and 72h post electroporation, and luciferase activity was measured as a surrogate for viral replication. All data shown are the average of three or more independent experiments. Error bar indicates the standard deviation of the mean and asterisk indicates significant differences, as determined by one-way ANOVA (*P<0.033, **P<0.002, ***P<0.001).

Thus, HCV mutant replication in Huh 7.5 Drosha KO cells was observed to be similar to their replication in Huh 7.5 miR-122 KO cells and suggest that the mutant viruses are capable of replicating in an environment devoid of miRNAs generated via the Drosha biogenesis pathway. The miRNAs found previously to support HCV replication, miR-504-3p, miR-574-5p and miR-1236-5p, miR-25-5p and miR-4730, are all generated by the canonical miRNA biogenesis, and thus we can conclude that viral replication in Huh 7.5 Drosha KO cells is independent of both miR-122 and miRNAs generated by the canonical Drosha dependent biogenesis, but we cannot rule out roles for mirtrons and other small RNAs generated independently from Drosha.

### HCV mutants replicating independently of miR-122 have greater translation efficiency compared to wild-type HCV

Studies from our lab and others have suggested that miR-122 binding to the HCV 5’ terminus stimulates virus translation, and that translation stimulation is the primary mechanism by which it promotes HCV propagation (12, 32). Further, *in silico* RNA structure predictions and biochemical analysis suggest that the 5’ UTR RNA may form a dynamic structure that modulates HCV IRES function. We and others have speculated that miR-122 annealing or 5’ UTR mutations that promote miR-122-independent HCV replication may shift the equilibrium to favor the formation of the active IRES **(Fig. 1**) (11, 12, 31). In addition, previous studies on miR-122-independent replication of bicistronic HCV construct suggested that altered translation regulation may enable miR-122-independent replication (33). Based on this, we hypothesize that viruses with 5’ UTR mutations that enable miR-122-independent replication would exhibit enhanced translation efficiency compared to the wild-type virus. To test our hypothesis, we have compared the translation efficiency of mutants and wild-type viral genomes in Huh 7.5 miR-122 KO cells (**Fig. 3A**). To focus on virus translation, we have used replication-defective virus mutants (GNN) in the translation assays, and a firefly luciferase (Fluc) messenger RNA (T7mRNA) was used as an internal control. The *Renilla* to Firefly luciferase ratio (Rluc:Fluc) was measured to calculate the translation efficiency and is expressed as percent luciferase expression relative to the translation by wild-type viral RNA **(Fig. 3A)**.

**FIG 3:**
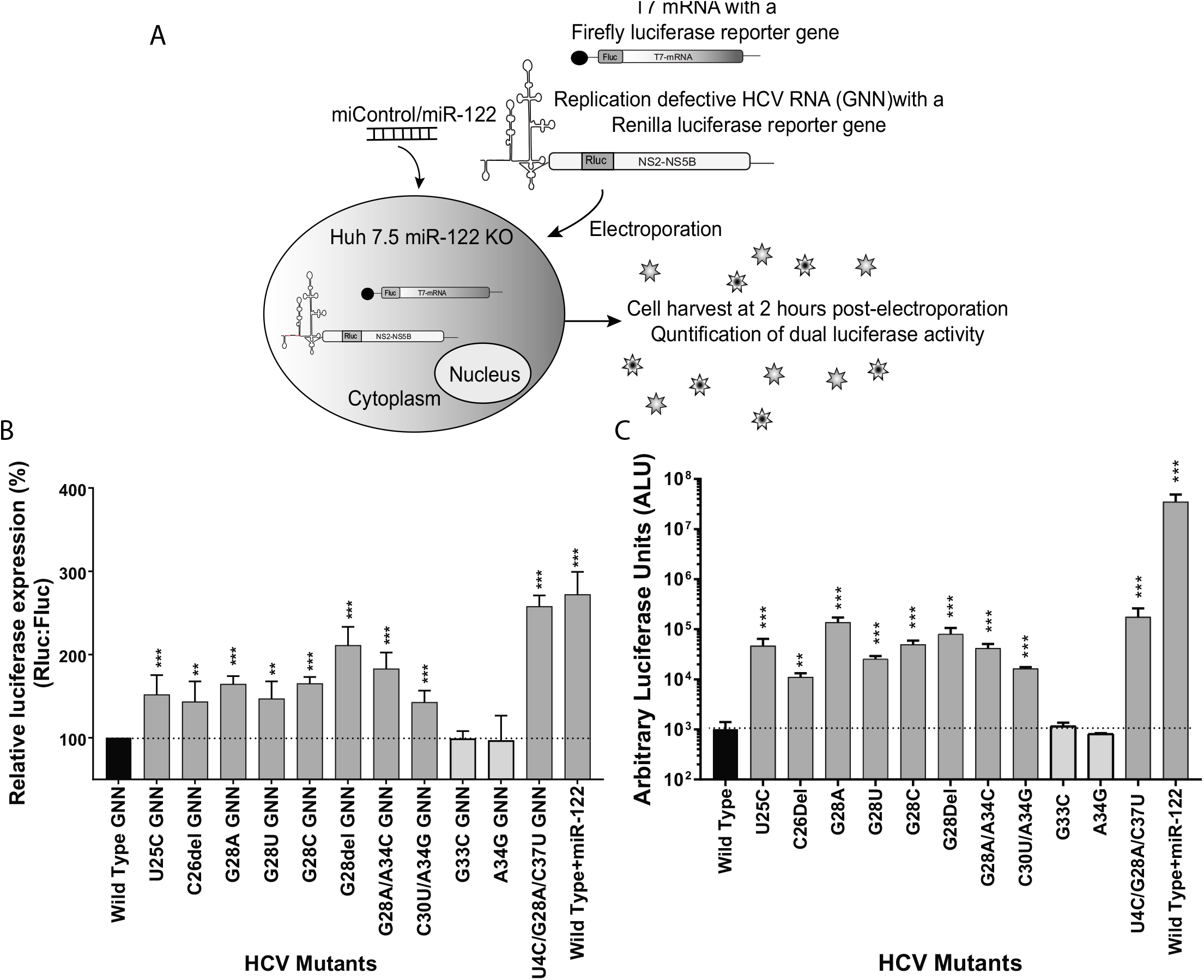
HCV mutants replicating independent of miR-122 have higher translation efficiency compared to wild-type HCV, and miR-122-independent translation potency correlates with miR-122-independent replication efficiency. (A) Illustration of the translation assay method. Replication defective *Renilla* reporter J6/JFH-1 (p7Rluc2a) HCV RNA, and a Firefly reporter mRNA were co-electroporated in Huh 7.5 miR-122 knockout cells. Cells were harvested 2 hours post electroporation and dual luciferase activity was measured. (B) Comparison of translation efficiency of HCV mutants and wild-type virus. (C) Replication of the mutant and wild-type HCV in Huh 7.5 miR-122 knockout cells. HCV RNA was electroporated into Huh 7.5 miR-122 knockout cells and cells were harvested 72 hours post-electroporation. *Renilla* luciferase was measured to analyze viral propagation. (B and C) In both translation and replication assay, wild-type virus is indicated in black whereas HCV mutants reported of replicating independently of miR-122 are indicated in grey, and HCV mutants that do not replicate independently of miR-122 are indicated in light grey. All data are presented as the average of three or more independent experiments. Error bar indicates the standard deviation of the mean and asterisk indicates significant differences. The significance was determined by using the Student’s t-test (*P<0.033, **P<0.002, ***P<0.001).

In support of our hypothesis, the translation assays showed that HCV mutants capable of replicating independently of miR-122 displayed higher translation efficiency compared to the wild-type virus and virus mutants incapable of miR-122-independent replication (**Fig. 3B**). The level of enhanced translation also correlated with the efficiency of miR-122-independent replication in Huh 7.5 miR-122 KO cells (**Fig. 3C**). For example, the mutant U4C/G28U/C37U exhibited the highest translation efficiency compared to other mutants and also the highest miR-122-independent replication efficiency, mutants with intermediate translation efficiency (G28A, G28del, and G28C) and U25C exhibited intermediate miR-122-independent replication, and mutants that exhibited translation lower than other mutants replicated poorly in the absence of miR-122 (C26del, G28U, and C30U/A34G,). Finally, the translation efficiency of mutants incapable of replicating independently of miR-122 (A34G, and G33C), were similar to that of wild-type virus. The result supports our hypothesis that enhanced translation efficiency by 5’ UTR mutant viruses allows HCV to replicate independently of miR-122 and suggests a role for miR-122 in viral translation regulation.

### The influence of other miR-122 binding site mutations on translation and miR-122-independent replication

Several reports have used viruses having mutations to the miR-122 binding sites to investigate the impact of miR-122 on virus replication (27, 39, 40). For example, we and others have used S1:p3, and S1:p4 (C26G, and U25A) mutants to study the impact of abolishing miR-122 annealing. In addition, other reports have used an array of miR-122 binding site mutants (S2 p1, p2, p3, p4, p5 to determine the impact of miR-122 annealing at each nucleotide (40), and on the ability of viruses to replicate and sometimes revert (41). However, these mutants have not been assessed in transient replication assays for their impact on miR-122-dependent and miR-122-independent replication. In light of more recent evidence that point mutations at miR-122 binding sites facilitating miR-122-independent replication of HCV, we wanted to revisit these mutants to assess whether any of them are capable of miR-122-independent replication (11, 34). To investigate whether mutations in the miR-122 binding sites altered virus translation or allow the virus to replicate in the absence of miR-122, we mutated each nucleotide of both the miR-122 binding sites individually to their complementary nucleotide and tested for virus replication in the absence of miR-122 in Huh 7.5 miR-122 KO cells. Altogether, we created 12 mutants; 6 for site 1, S1p2 to S1p7 and 6 for site 2, and S2p2 to S2p7 (**Fig. 4A**). Replication of these mutants was also tested in the presence of miR-122 to verify their replication efficiency when one miR-122 site is bound.

**FIG 4:**
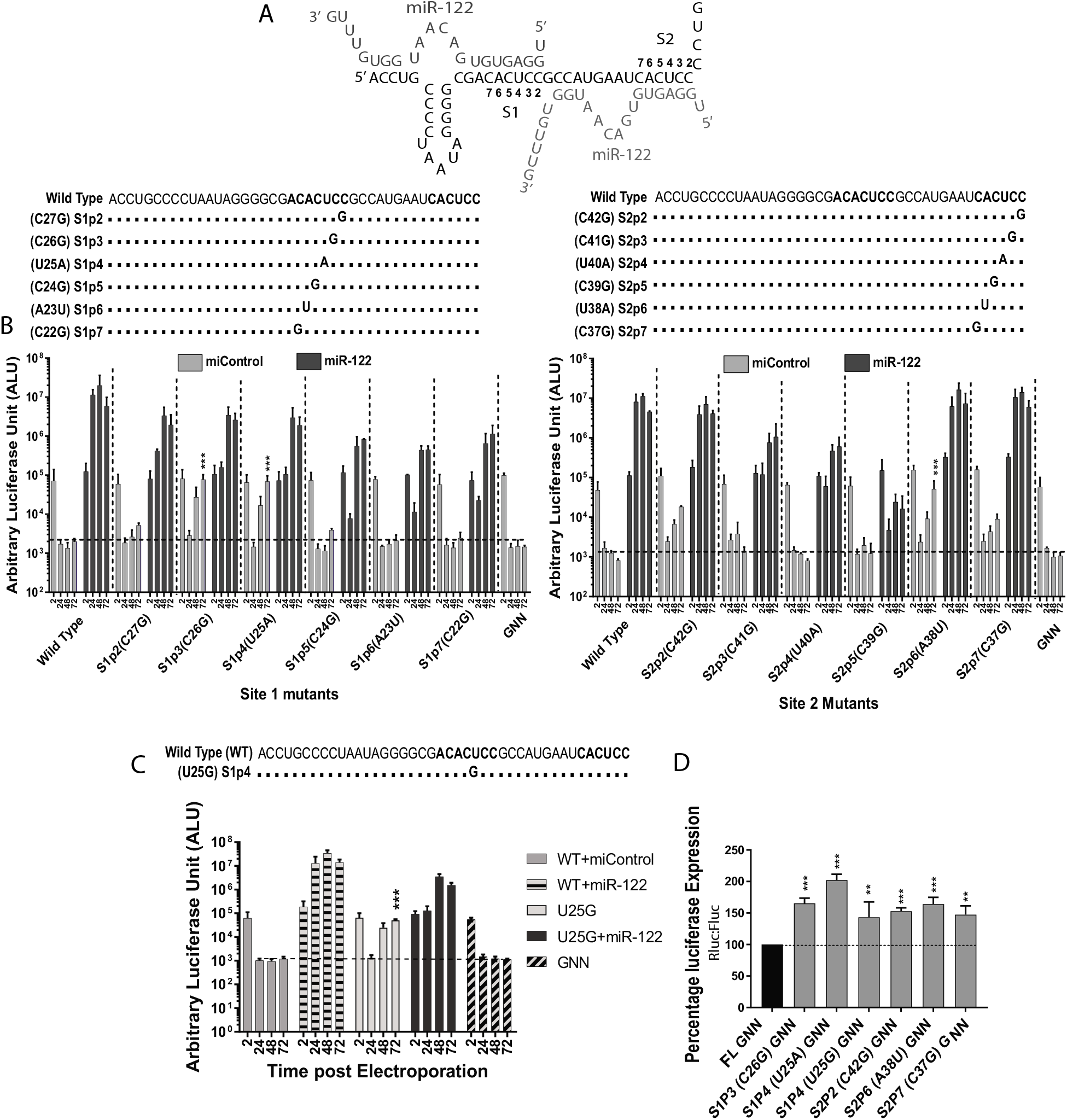
Some mutations to nucleotides within miR-122 binding sites 1 and 2 allow viral propagation in the absence of miR-122. (A) The HCV 5’ UTR and the location and name of mutants viruses with sequence changes within miR-122 binding sites 1 and 2. ‘S’ stands for miR-122 binding site (1 or 2) and ‘p’ stands for the position in the binding site (1 to 7). (B) Replication of Site 1 and Site 2 mutant HCV J6/JFH-1 (p7Rluc2a) RNA in Huh 7.5 miR-122 knockout cells with a control miRNA (miControl) or with miR-122. *Renilla* luciferase was assessed at 2h, 24h, 48h, and 72h post electroporation as an indicator of viral propagation. The Dark grey bars and the light grey represents miR-122-dependent and independent replication, respectively. (C) Replication of HCV J6/JFH-1 (p7Rluc2a) U25G mutant RNA in Huh7.5 miR-122 knockout cells with or without miR-122. miR-122 independent replication of HCV U25G RNA was compared to the wild-type HCV RNA with miControl as a control. (D) Comparison of translation of site 1 and site 2 HCV J6/JFH-1 (p7Rluc2a) mutant RNA with that of J6/JFH-1 (p7Rluc2a) wild-type replication defective mutant. All data are presented as the average of three or more independent experiments. Error bar indicates the standard deviation of the mean and asterisk indicates significant differences. The significance was determined by using one-way ANOVA (*P<0.0332, **P<0.002, ***P<0.001).

All of the mutants, except for S2p5 (C39G), could replicate to wild-type levels in the presence of miR-122 (**Fig. 4B**). The replication of S2p5 was abolished even in the presence of miR-122 suggesting that C39G mutation has a severe effect on viral replication fitness. In addition, some mutants could replicate independently of miR-122, and miR-122-independent replication correlated with translation efficiency. Specifically, three mutants S1p3, S1p4, and S2p6 (U26G, U25A, and A38U) displayed miR-122-independent replication and significantly enhanced translation efficiency in Huh 7.5 miR-122-KO cells (**Fig. 4D**). The translation efficiency of S2p2 and S2p7 (C42G and C37G) was also found to be significantly higher than wild-type virus (**Fig. 4D**), and while these mutants showed evidence of miR-122-independent replication, luciferase expression levels were not statistically different than that from wild-type virus (**Fig. 4B**). Thus, our result supports our hypothesis suggesting that HCV mutants with an enhanced translation ability can also replicate independently of miR-122.

### Any other nucleotides at positions C25 and G28 enhance translation and support miR-122-independent replication

In a previous report from our lab, we have shown that any nucleotide except G at position 28 of the HCV 5’ UTR, including a deletion (the ‘G28 mutants’) can support miR-122-independent replication (11). Here we show that the G28 mutants supporting miR-122-independent replication also translated more efficiently than the wild-type virus and their translation efficiency correlated with their replication efficiency. Similarly, while studying S1 and S2 mutants, we have found that virus genomes having any nucleotide except U at position 25 can also replicate independently of miR-122. U25C has been reported previously to replicate independently of miR-122 (11, 34), and we show that U25A and U25G (**Figs 4B and C)** can also replicate independently of miR-122. Translation analysis of the U25 series of mutants also shows higher translation efficiency compared to the wild-type virus (**Fig. 4D**) and further supports our hypothesis that enhanced IRES translation efficiency enables miR-122-independent replication.

### HCV with miR-122 site 2 mutations allow the virus to replicate independently of miR-122

Mutation of miR-122 binding site 2 from CACUCC to GGCGUG (HCV-S2-GGCGUG) was also reported of enabling efficient miR-122-independent replication (37). In this mutant, all positions in S2 except for S2p5 were mutated, suggested that site 2 binding of miR-122 is dispensable for HCV propagation and that its mutation can rescue miR-122-independent replication. In support of our model that miR-122-independent replication is facilitated by enhanced translation, we found HCV-S2-GGCGUG to exhibit enhanced translation ability compared with the virus having a wild-type 5’ UTR (**Fig. 5C**). To test the role of abolishing miR-122 binding site 2, we have mutated each nucleotide of the miR-122 binding site 2 to its complementary nucleotide (CACUCC to GUGAGG) (HCV-S2C) and tested its replication and translation in Huh 7.5 miR-122 binding KO cells. HCV-S2C also replicated independently of miR-122 (**Fig. 5A**) but about 10-fold less efficiently than HCV-S2 (**Fig. 5B**), and the translation ability of the mutants correlated with their miR-122-independent replication ability (**Fig. 5C**). Thus, mutations within the S2 region modulate the ability of the 5’ UTR to regulate virus translation and virus dependence on miR-122.

**FIG 5:**
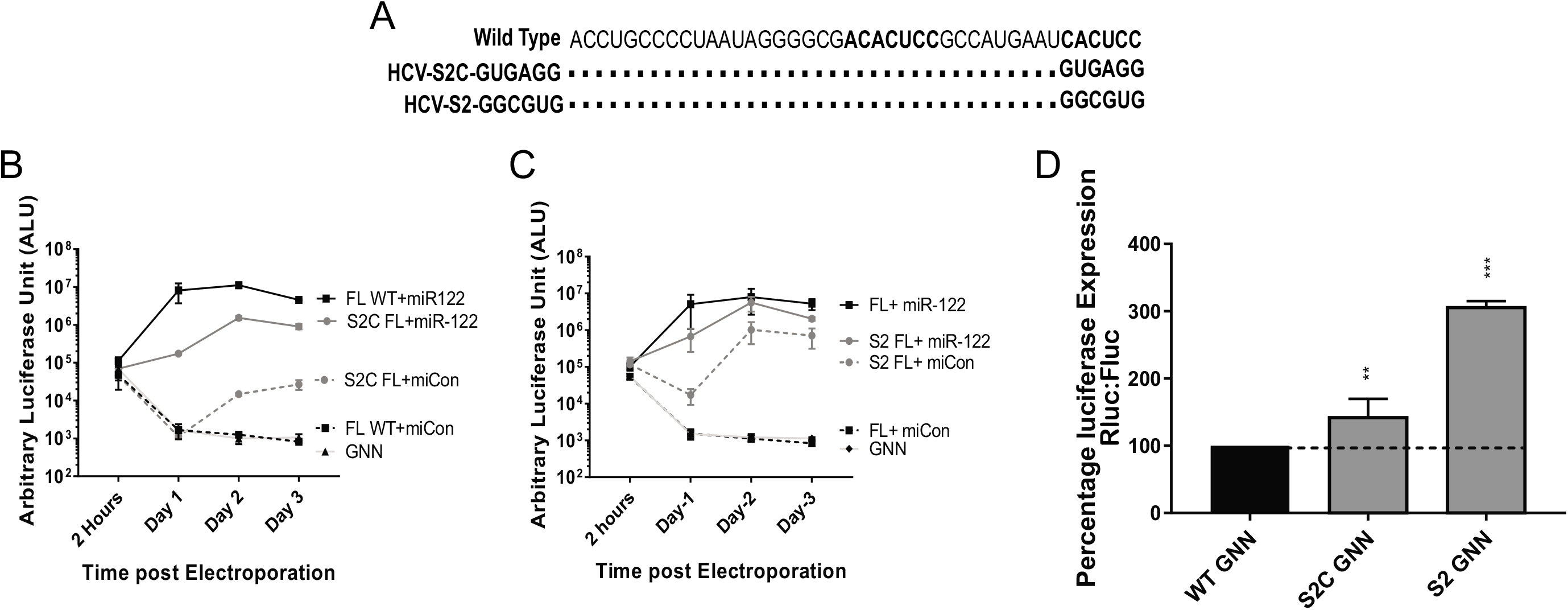
Translation and replication assays of miR-122 binding site 2 HCV mutants that replicate independently of miR-122: (A) 5’ UTR sequence (1-42 nucleotide) of miR-122 binding site 2 mutants. Replication of J6/JFH-1 (p7Rluc2a) HCV-S2C-GUGAGG (B) and HCV-S2-GUCGUG (C) in Huh 7.5 miR-122 knockout cells with or without miR-122. Wild-type and mutant HCV J6/JFH-1 (p7Rluc2a) RNA was electroporated with miControl or miR-122 and cell were harvested at 2h, 24h, 48h, and 72h, and luciferase activity was measured as an indicator of viral propagation. (D) Translation of HCV miR-122 binding site 2 mutants compared to the wild-type replication defective mutants. All data shown are the average of three or more independent experiments and the error bar indicates the standard deviation of the mean and an asterisk indicates significant differences. The significance was determined by using one-way ANOVA (*P<0.033, **P<0.002, ***P<0.001)

### miR-122 binding rescues viral translation of translationally impaired SLII mutation

A proposed mechanism by which miR-122 promotes HCV translation is that its annealing promotes the formation of the translationally active SLII RNA structure and that in the absence of miR-122, the 5’ UTR has a greater propensity to fold into a translationally inactive structure called SLII^Alt^. In addition, we and others speculate that mutations in the 5’ UTR that stimulate translation and promote miR-122-independent HCV replication favour the translationally active structure even in the absence of miR-122 (**Fig. 1**), but this remains to be confirmed biophysically(11, 12). In an attempt to link translation efficiency and RNA secondary structure using virus genetics, we generated a mutant virus predicted to be less capable of forming SLII^Alt^ by modifying the sequence of SLII instead of the 5’ terminal region. We chose to mutate A98C because it is predicted to pair with nucleotide U25 within the SLII^Alt^ structure, similar to the U25C mutation, and mutation of A at position 98 is predicted to weakens the SLII^Alt^ structure and induce the translationally active SLII structure. However, mutation of A98 to C impaired both translation and replication (**Fig. 6A**). Viral luciferase expression levels at the 2-hour time point were significantly lower than from wild-type virus suggesting the negative effect of SLII mutation in the viral translation. This RNA virus was also incapable of miR-122-independent replication. Supplementation with miR-122 rescued both translation (2hour time point) and replication of A98C but replication was significantly lower compared to the wild-type RNA with miR-122. While these data do not provide insight into RNA structures, they support our central hypothesis that viral translation is correlated with overall viral replication and that miR-122 binding supports viral propagation by promoting viral translation (**Fig. 6B**). A role for miR-122 in promoting translation was further supported by a mutant having a 19-nucleotide insertion into SLII that was generated during A98C mutagenesis (Ins Mutant). The insertion is predicted to completely abolish SLII formation (**Fig. S1**) and, as expected, translation of luciferase from this construct was ablated. However, miR-122 binding rescued viral translation of the Ins mutant suggesting miR-122 can rescue translation of RNAs predicted to have severely altered SLII structures.

**FIG 6:**
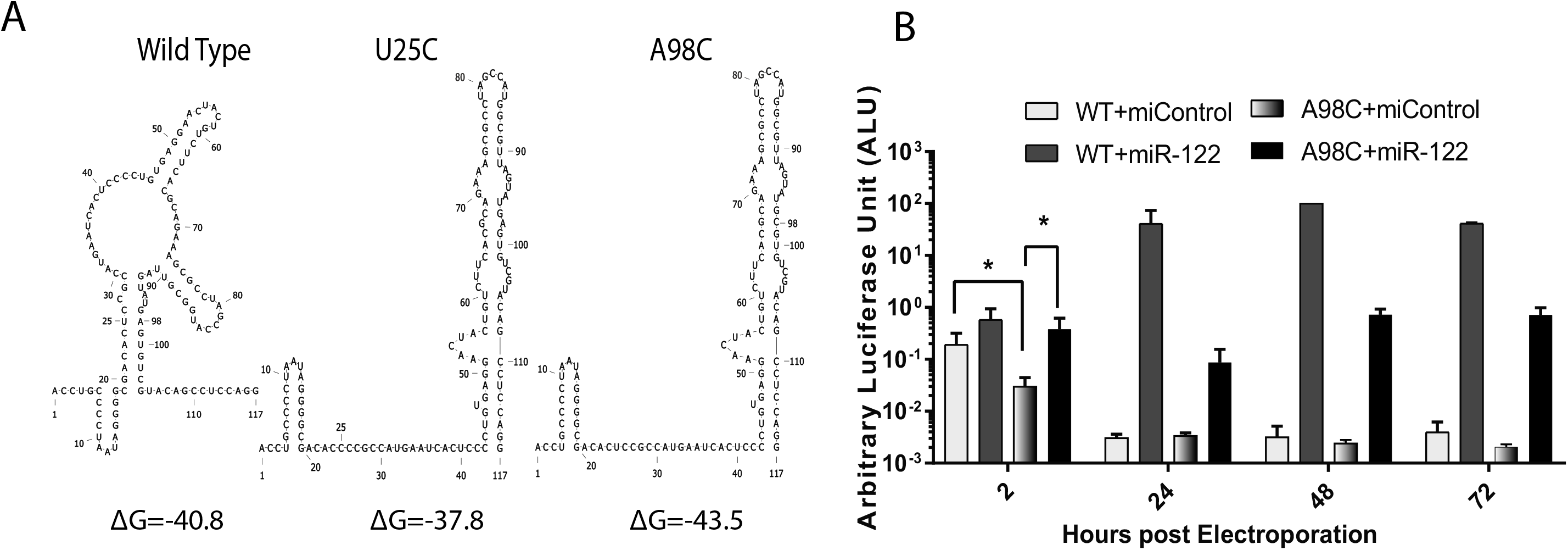
A mutation in SLII that is predicted to modify the 5’ UTR RNA structure does not confer miR-122-independent replication (A) Structure prediction of 1-117 nucleotide of J6/JFH-1 (p7Rluc2a) wild-type HCV, U25C HCV, and A98C HCV. The first favorable lowest free energy structure is represented. Wildt-type HCV is predicted to form the non-canonical SLII^Alt^ structure along with SLI whereas both U25C and A98C HCV are predicted to form the canonical structure with SLI and SLII. (B) Replication of wild-type and A98C HCV mutant in the presence of control miRNA or miR-122 in Huh7.5 miR-122 knockout cells. Error bar indicates the standard deviation of the mean and asterisk indicates significant differences. The significance was determined by using Student’s t-test (*P<0.033, **P<0.002, ***P<0.001).

### Enhanced translation and genome stability can completely rescue HCV propagation in the absence of miR-122

Roles for miR-122 in the stimulation of HCV translation and stabilizing the genomic RNA have been reported but the relative contribution of each role to overall HCV propagation is still unknown. In this study, we have correlated enhanced translation stimulation with the ability of mutant viral RNAs to replicate independently of miR-122, but mutation-induced miR-122-independent replication is still 10-fold lower than miR-122-dependent replication. We hypothesize that lower miR-122-independent replication efficiency in the absence of miR-122 was due to lack of miR-122-annealing based genome stability, and that genome stabilization by other means will enhance miR-122-independent replication and reveal the relative impact of miR-122 in genome stabilization on overall HCV propagation. To test this hypothesis, we assessed the ability of the knockdown of RNA degrading enzymes in rescuing HCV replication in the absence of miR-122. Previously, miR-122-independent replication of HCV bicistronic replicon RNA had been rescued by knockdown of the cellular pyrophosphatases DOM3Z and DUSP11 and the cellular exonuclease XRN1 (28, 42). To assess the impact of genome stabilization on miR-122-independent replication of 5’ UTR mutants, we assessed rescued miR-122-independent replication of wild-type and mutant HCV (U4C/G28A/C37U, and U25C) (**Fig. 7A**) by knockdown of XRN1, DOM3Z, and DUSP11. Our result showed that miR-122-independent replication of U4C/G28A/C37U and U25C was similar to their miR-122-dependent levels in XRN1, DOM3Z, and DUSP11 knockdown environment (**Fig.7B**, Mutant HCV+miControl+siDOM3Z, siXRN1, siDUSP11 vs Mutant HCV+miR-122+siControl). miR-122-independent replication of the wild-type virus was also enhanced by knockdown of XRN1, DOM3Z, and DUSP11, but did not reach miR-122-dependent levels (**Fig. 7B**, wild-type HCV+miControl+ siXRN1, siDUSP11, siDOM3Z vs wild-type HCV+miR-122+siControl). From this experiment, we suggest that enhanced translation derived from 5’ terminal point mutations can completely rescue viral replication to miR-122-dependent level in the absence of RNA degrading enzymes and thus compensate for the role of miR-122 in the viral life cycle. The experiment also suggests that apart from translation, miR-122 also plays a role in stabilizing the viral genome by protecting it from enzyme degradation.

**FIG 7:**
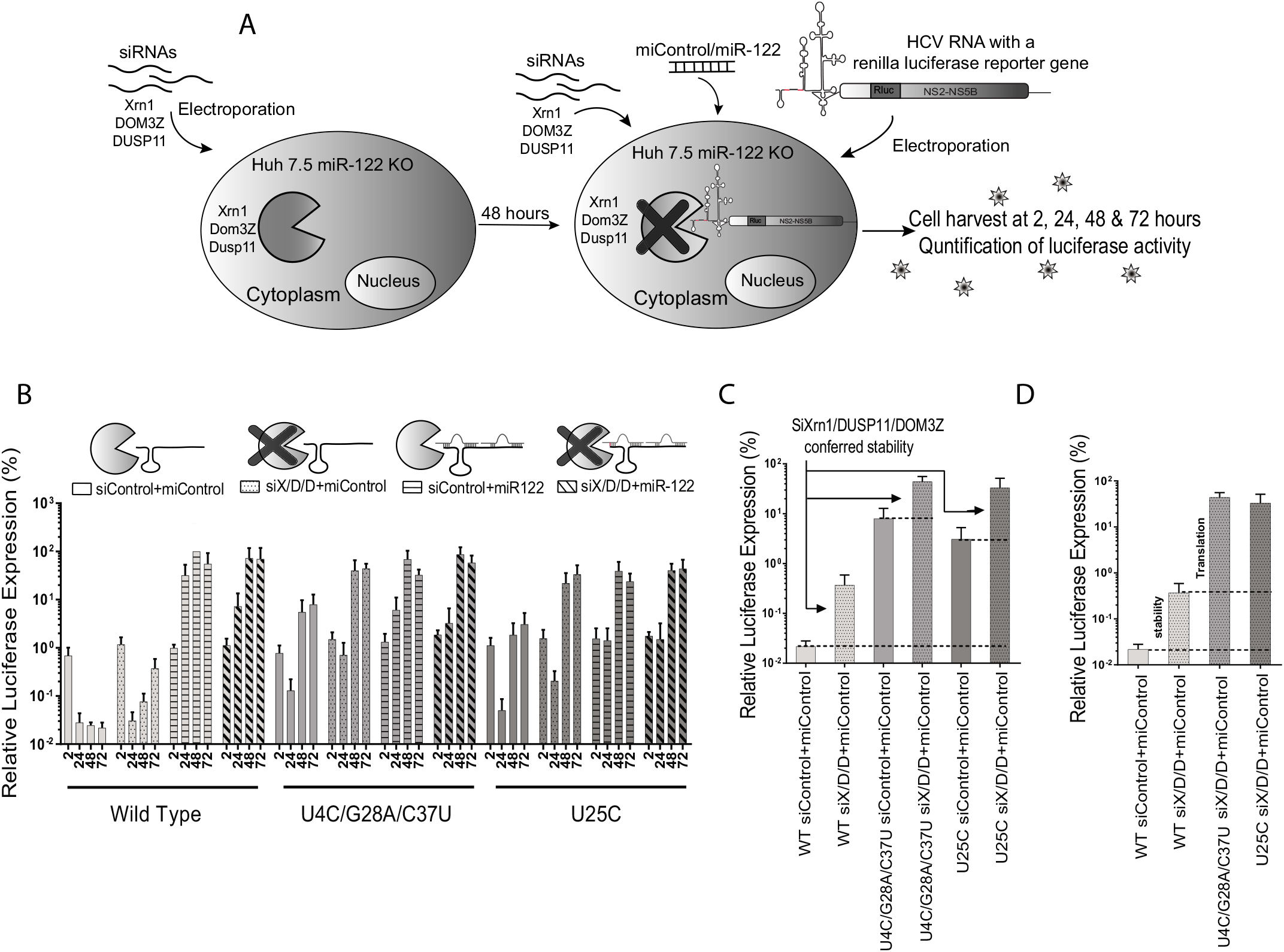
Viral genome stabilization rescues miR-122-independent replication of 5’ UTR mutants to wild-type HCV levels. (A) Graphical representation of siRNA induced XRN1, DUSP11, and DOM3Z knockdown and HCV replication analysis in Huh 7.5 miR-122 knockout cells. Huh 7.5 miR-122 knockout cells were electroporated with XRN1, DUSP11, and DOM3Z specific siRNAs and 48 hours after the first electroporation, viral RNAs along with siRNAs and miRNAs were co-electroporated into the preknockdown Huh 7.5 miR-122 knockout cells. Cells were harvested at 2h, 24h, 48h, and 72h post second electroporation, and luciferase activity was measured as an indicator of viral propagation. (B) Replication of J6/JFH-1 (p7Rluc2a) wild-type HCV, U4C/G28A/C37U HCV, and U25C HCV RNA in Huh 7.5 miR-122 knockout cells. The solid bars represents HCV replication without miR-122 or siRNA knockdown, dotted bars represent HCV replication without miR-122 and with siRNA knockdown, bars with horizontal line represents HCV replication with miR-122 and without siRNA knockdown, and bars with slanted lines represents HCV replication with miR-122 and with siRNA knockdown. (C) The siXRN1, siDUSP11, and siDOM3Z conferred stability was determined for wild-type, U4C/G28A/C37U, and U25C HCV RNA by comparing replication without knockdown (siControl) vs replication with knockdown (siS/D/D). (D) The data shown in C was used to determine the contribution of stability to miR-122-independent replication by comparing replication of wild-type HCV RNA in miR-122 knockout cells with (siX/D/D) and without (siControl) knockdown of XRN1, DUSP11, and DOM3Z. Similary, the contribution of mutation induced translation to miR-122-independent replication was determined by comparing replication of stabilized wild-type HCV RNA (WT siX/D/D) with stabilized mutant RNAs (U4C/G28A/C37U siX/D/D) and (U25C siX/D/D).

We also used this assay to determine the relative impact of miR-122 on viral translation and genome stability by comparing the replication of wild-type virus and mutant viruses with and without knockdown of XRN1, DOM3Z, and DUSP11. Knockdown of XRN1, DOM3Z, and DUSP11 increased miR-122-independent luciferase expression of wild-type, U4C/G28A/C37U, and U25C viruses by almost 10-fold compared to luciferase levels without knockdown (**Fig.7C**). Thus, we conclude that miR-122-induced RNA stabilization accounts for about a 10-fold increase in replication. In addition, we assessed the relative impact of translation stimulation on HCV replication by comparing luciferase levels from wild-type vs mutants in DOM3Z, XRN1, and DUSP11 knockdown cells (**Fig.7D**). In this experiment, we removed the need for miR-122 in viral genome stabilization by knocking down DOM3Z, XRN1, and DUSP11, thus allowing the experiment to isolate the impact of the translation enhancing mutations on virus replication. In this case, the mutant viruses replicated about 100-fold vs wild-type virus and we conclude that miR-122-induced translation stimulation accounts for about a 100-fold increase in HCV replication. Altogether, our results indicated that translation stimulation is the primary mechanism by which miR-122 promotes HCV propagation, and that genome stabilization is an important but less potent mechanism.

We further support the notion that genome translation simulation alone is not sufficient to promote miR-122-independent HCV replication. We attempted to rescue HCV replication to miR-122-dependent levels by enhancing translation using two methods, 5’ UTR mutations and the addition of an EMCV IRES. In our previous work, we showed that miR-122-independent replication was also promoted by altered viral protein translation regulation by an Encephalomyocarditis virus (EMCV) IRES in a subgenomic HCV (SGR JFH-1 Fluc). In this experiment, we tested miR-122-independent replication of a bicistronic sub-genomic HCV construct that also had 5’ UTR mutations (U4C/G28A/C37U SGR JFH-1 Fluc) that promote miR-122-independent replication to test if we could further enhance miR-122-independent replication (**Fig. 8**). The U4C/G28A/C37U SGR JFH-1 Fluc replicated better than SGR JFH-1 Fluc in the absence of miR-122; however, even with the presence of two translation enhancing genetic features the miR-122-independent replication of U4C/G28A/C37U SGR JFH-1 Fluc was not equal to the miR-122 dependent level (**Fig. 8**). Assessing the assumption that translation might be the major contributor of miR-122’s function in HCV replication, data from the above experiment suggest that miR-122’s function in genome stability might be contributing a minor part in HCV propagation but it is still essential for efficient HCV propagation.

**FIG 8:**
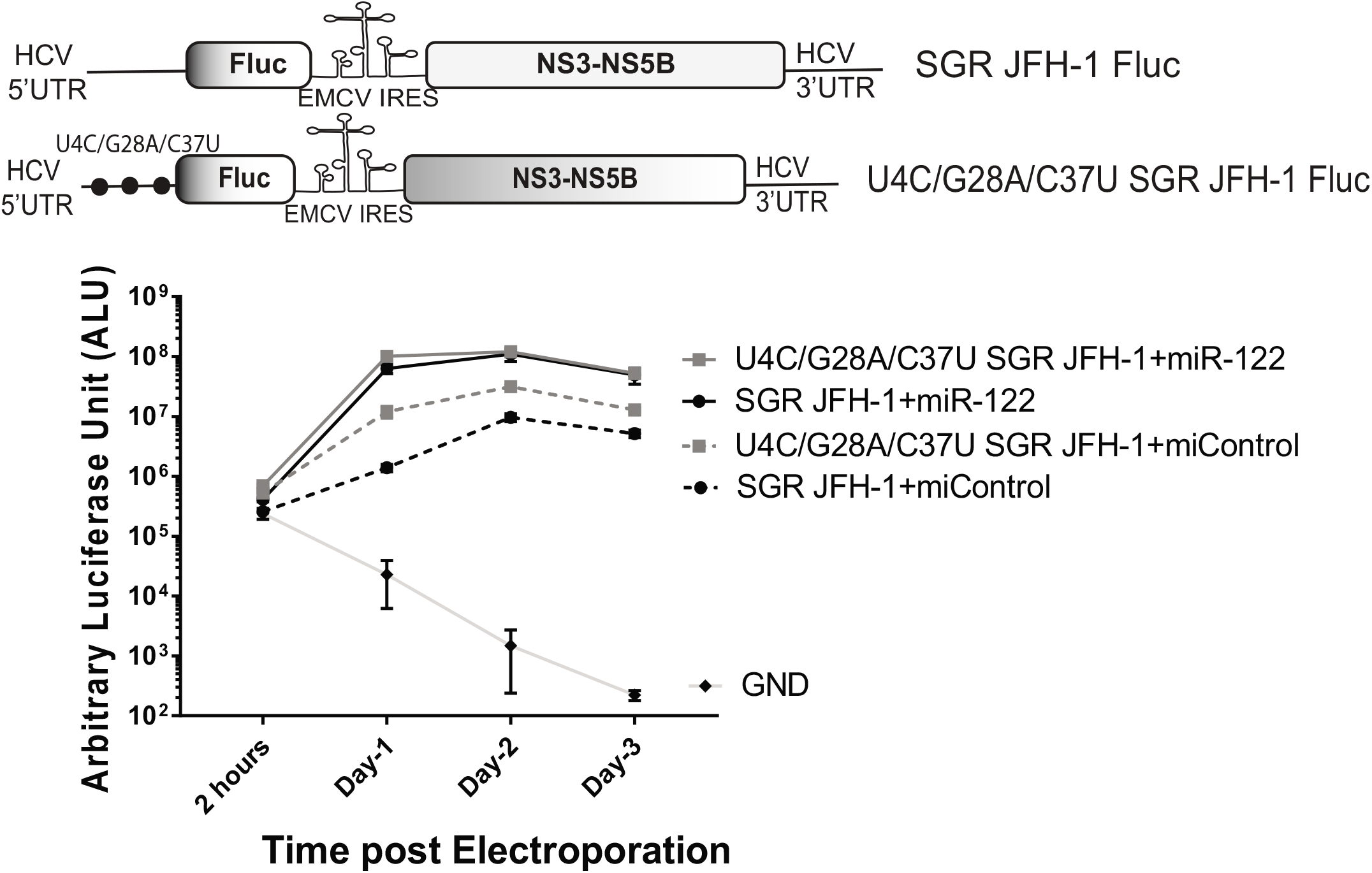
Enhanced translation alone cannot rescue HCV propagation to miR-122 dependent level. A wild-type bicistronic subgenomic HCV (SGR JFH-1 Fluc wild-type) and a triple mutant bicistronic subgenomic HCV (U4C/G28A/C37U SGR JFH-1 Fluc) were electroporated into Huh 7.5 miR-122 knockout cells with or without miR-122, and cells were harvested at 2h, 24h, 48h, and 72h time points to analyse the Firefly luciferase activity, which is a measure of viral replication inside the cell. Black lines represent wild-type SGR JFH-1 Fluc and grey lines represent U4C/G28A/C37U SGR JFH-1 Fluc. GND is the replication defective negative control.

## DISCUSSION

In this study, we have investigated miR-122-independent replication of HCV in an effort to understand the role of miR-122 in promoting the HCV life cycle and have used several virus mutants having 5’ UTR mutations reported by us and others that support miR-122-independent replication. However, A recent report suggests that HCV replication can be supported by other miRNAs (miR-504-3p, miR-574-5p, and miR-1236-5p) in a miR-122-like manner and by miR-25-5p and miR-4730 in a non-miR-122 like manner (39). To investigate the possible effects of these miRNAs on miR-122-independent HCV replication we assessed replication of several 5’ UTR mutants in Huh 7.5 Drosha KO cells. Drosha is required for the canonical miRNA biogenesis pathway and thus Huh 7.5 Drosha KO cells do not express the above mentioned and most other miRNAs, including miR-122. We found that the mutant viruses replicating in the absence of miR-122 could also replicate in Huh 7.5 Drosha KO cells suggesting that other miRNAs made by the canonical miRNA biogenesis pathway are not supporting miR-122-independent HCV replication in our assays. However, we cannot exclude the possibility that miR-122-independent replication might be supported by small RNAs generated by non-canonical pathways such as mirtrons, snoRNA derived miRNAs and tRNA derived miRNAs (43, 44).

Our analyses of miR-122-independent replication suggest that translation stimulation is the primary role for miR-122 in promoting the virus life cycle, and that genome stabilization has a less potent impact but is still important. In this study, we have explored the translation efficiency of several mutants that replicate independently of miR-122 and found that all mutant viruses capable of miR-122-independent replication also had enhanced translation abilities, and that the translation strength of the mutant genomes correlated with miR-122-independent replication abilities (**Fig. 3**). This suggests that translation stimulation by miR-122 contributes to HCV replication promotion, and mutants with enhanced translation efficiency can compensate for the lack of miR-122. This supports our previous finding that altered translation regulation may compensate for miR-122’s function in viral propagation (33) based on miR-122-independent replication of bicistronic JFH-1 SGR RNAs in which viral protein expression is driven by the EMCV IRES instead of the HCV IRES. Translation stimulation was proposed as a mechanism by which miR-122 promotes the HCV lifecycle several years ago, but the small stimulation of translation observed, approximately 2-fold, appeared insufficient to account for the dramatic effect on virus replication. However, a recent study on translation and replication dynamics of picornavirus (positive-strand RNA virus) suggest that the efficiency of initial viral translation and production of a sufficient amount of protein to initiate genome replication is important to establish an infection, and, in cases where there is an insufficient translation and failed replication, the virus reinitiate translation for another attempt to establish an infection (45). The study also suggests that viral translation occurs at multiples phases of the viral life cycle. Hence, suggests that a small translation stimulation by miR-122 detected in translation assays could have a significant and compounding role throughout the viral life cycle.

A second proposed mechanism by which miR-122 promotes the HCV life cycle is by stabilizing the viral genome, and our results indicate that miR-122-induced genome stabilization has a less potent impact than translation stimulation but is still important for efficient virus propagation. While we were not able to directly compare the stability of the mutant viral RNA in cells, we analyzed the contribution of genome stability to mutant RNA replication by analyzing it in the absence of RNA degradation proteins. We found that miR-122-independent replication of mutant viruses was rescued to miR-122-dependent levels by knockdown of exonuclease XRN1, and pyrophosphatases DOM3Z and DUSP11 suggesting that genome stabilization allows the mutants with enhanced translation to replicate to miR-122 dependent level. This supports the role of miR-122 in both translation stimulation and genome stabilization and that wild-type levels of miR-122-independent replication can be achieved by providing these roles using alternative methods. Further, we were also able to determine the relative contributions of translation stimulation and genome stabilization to 100-fold and 10-fold respectively. Although we suggest translation as a major contributor to miR-122’s function in viral propagation, we have also shown that miR-122 induced genome stability is a minor but essential function in HCV propagation.

In addition, we and others have reported that mutations on the 5’ UTR that promote miR-122-independent replication also promote the favorable secondary structure of the active HCV IRES (**Fig. 1),** and confirmed the high translation efficiency of G28A, a mutant capable of miR-122-independent replication (12).

We and others have proposed a model that binding of miR-122 to the 5’ UTR induces the formation of the translationally active RNA secondary structure (11, 12, 31, 32). The proposed model suggests the 5’ UTR RNA forms one or more translationally unfavorable structures (SLII^Alt^) in the absence of miR-122 and that miR-122 annealing modifies the thermodynamics to favor the formation of the translationally active SLII structure. Based on extensive structure-function analyses and X-ray crystallography, nucleotides 40 to 372 of the HCV RNA forms an IRES structure in solution (13, 46) and the model proposes that the 5’ terminal nucleotides 1-42, including the miR-122 binding sites modulates the structure of the IRES. However, biophysical or structural analyses are lacking and X-ray crystallography experiments were done using RNAs that lacked the 5’ terminal region containing the miR-122 binding sites. We propose that 5’ UTR point mutants that promote miR-122-independent replication also shift the equilibrium to favor the formation of SLII, even in the absence of miR-122 (**Fig. 1**). In support of this hypothesis, the mutants we found to have enhanced translation are also predicted to favour the formation of the canonical SLII prediction structures in the absence of miR-122 (11, 31) (**Fig. 3**) (**Fig. S2 and S3**) By contrast; however, some of the mutants that replicated independently from miR-122 were not predicted to form the translationally active SLII structure. Specifically, Site 2 mutants, HCV-S2-GGCGUG and HCV-S2C-GUGAGG were predicted to form a secondary structure in which the 5’ terminal sequences folds to generate a single long stem loop (**Fig. S4**), and another site 2 mutant S2p6 (U38A) was predicted to form other RNA structures. Further, structure prediction analysis has also suggested other exceptions where mutant sequences are predicted to form the canonical structure but do not exhibit miR-122-independent replication or enhanced translation (**FIG. S3)**, and thus suggest the mechanism might be more complex. In addition, the model does not consider the roles of RNA binding proteins recruited or displaced by miR-122 which have been proposed to also modulate 5’ UTR structures (31). Thus, confirmation or refinement of the model will require biophysical analyses of the RNA structures generated by these mutants and covariance of RNA structures with virus replication phenotypes. The dynamic nature of the RNAs will also require a method such as Cyro-EM to resolve the different structures it may form with or without miR-122 annealing.

In the process of studying miR-122-independent replication of these mutants, we have identified a few 5’ UTR mutants that, despite being predicted to form translation favourable RNA structures, do not replicate at all, even in the presence of miR-122. We speculate that these mutations affect the structure and function of the complement strand of the 5’ UTR during genome replication. For example, A site 2 mutant S2p5 (C39G) (**Fig. 4B**) could not replicate, even in the presence of miR-122, despite the prediction that the 5’ UTR should form the canonical SLII structure (data not shown). However, this mutation is predicted to alter the structure of the complementary strand (**Fig. S4**). A study by Friebe *et al*. showed that the RNA secondary structure of the negative strand affects viral RNA replication. Thus, we suggest that mutations in the 5’ UTR of the viral genome may affect miR-122 annealing and the secondary structure of both strands and this could limit the tolerance of the 5’ UTR to point mutations. Thus, we suggest that miR-122-independent replication of HCV is dictated by the multifunctional nature of the 5 UTR and its complementary strand.

It is also interesting that while numerous HCV mutants capable of miR-122-independent replication have been isolated in the lab (**Fig. 3B**), few are found in patients’ samples. Only the mutant G28A was found in hepatic and extrahepatic tissue of patients as well as in cell culture supporting miR-122-independent replication and suggests evolutionary pressure to retain a dependence on miR-122, perhaps to restrict HCV replication to the liver. In addition, nonprimate hepaciviruses have at least one conserved miR-122 binding site on their genome suggest an evolutionary relationship to retain dependence on miR-122 (37, 47).

In conclusion, we have established that viral variants that replicate independently of miR-122 also display enhanced translation. We have also demonstrated that viral replication in the absence of miR-122 can be completely rescued when both translation and genome stability is addressed by other means. Finally, we show evidence that translation stimulation is a major function of miR-122 induced promotion of HCV whereas genome stability has a secondary function. Based on our study we propose that HCV mutants capable of miR-122-independent replication, are selected based on enhanced translation but must also retain replication fitness.

## MATERIALS AND METHODS

### Cells

Human hepatoma cell lines Huh 7.5 (48) and Huh 7.5 Drosha KO cells were kind gifts from Dr. C M Rice (49). Huh-7.5 miR-122 KO cell line was gifted by Dr. Matthew Evans (34). All the cells were cultured in Dulbecco’s modified Eagle medium (DMEM) supplemented with 10% fetal bovine serum, 0.1 nM non-essential amino acids (Wisent, Montreal, Canada), and 100 μg/mL Pen/Strep (Invitrogen, Burlington ON, Canada).

### Plasmids

Plasmids pJ6/JFH-1 Rluc (p7-Rluc2a) encoding full-length viral sequences derived from the J6 (structural proteins) and JFH-1 (non-structural proteins) isolates of HCV, and a *Renilla* luciferase reporter (Rluc) gene, directly downstream of the p7 gene was provided by Dr. C M Rice (herein called pJ6/JFH-1 RLuc) (50). The replication-defective mutant has a GAA-GNN mutation in the viral polymerase active site, pJ6/JFH-1 Rluc GNN. All site 1, site 2, and other 5’ UTR mutants were created by site-directed mutagenesis PCR of PBSKS+ vector with a cloned fragment of pJ6/JFH-1 Rluc plasmid from *EcoR*I to *Kpn*I. After confirming the mutation(s) by sequencing, digested *EcoR*I to *Kpn*I fragment from PBSKS+ plasmid was swapped into pJ6/JFH-1 Rluc WT plasmid. To generate mutant HCV variants with mutations in the 5’ UTR were cloned into the pJ6/JFH-1 Rluc by swapping *EcoR*I to *Kpn*I fragments. Replication defective mutants of all the viral variants were created by inserting the 5’ UTR mutations *EcoR*I to *Kpn*I fragment into pJ6/JFH-1 Rluc GNN. A pT7 plasmid encoding a messenger RNA with a firefly reporter gene was used to generate control mRNA for translation assays (Promega, Madison, USA), herein known as pT7 Fluc. Plasmid pSGR JFH-1 Fluc contains bicistronic JFH-1-derived subgenomic replicon (SGR) cDNAs with a firefly luciferase reporter (51). pSGR JFH-1 Fluc U4C/G28A/C37U plasmid was cloned by swapping *EcoR*I-*Age*I fragment from pJ6/JFH-1 Rluc (p7-Rluc2a) plasmid containing U4C/G28A/C37U mutation to pSGR JFH-1 Fluc. Plasmid pSGR JFH-1 Fluc was a gift from Dr. Takaji Wakita.

### microRNA and siRNA sequences

The sequence of synthetic miR-122 guide strand was UGGAGUGUGACAAUGGUGUUUGU, and the passenger strand was AAACGCCAUUAUCACACUAAAUA. miControl is a mutated version of miR-122 in which sites 2–8 on the guide strand were converted to their complement; guide strand UAAUCACAGACAAUGGUGUUUGU and the passenger strand AAACGCCAU UAUCUGUGAGGAUA (27). siXRN1 (si29015; 5′-GAG AGU AUA UUG ACU AUG Att-3′), siControl (5′-GAA GGU CAC UCA GCU AAU CAC ttc-3′) (27). siDOM3Z SMARTpool siGENOME were obtained from Dharmacon (Lafayette, CO, USA), siDUSP11 (AM16708) was purchased from Thermo Fisher Scientific (Waltham, MA, USA).

### Invitro transcription

pJ6/JFH-1 Rluc plasmids were linearized by digesting with *Xba*I and Mungbean nuclease and *invitro* transcription was performed using MEGA Script T7 High Yield Transcription Kit (Life Technologies, Burlington, Canada). To generate T7 Fluc mRNA, pT7 Fluc plasmid was linearized with *Xmn*I and mMessage mMachine mRNA synthesis kit (Life Technologies, Burlington, Canada) was used for *invitro* transcription (All the transcription process was performed using suggested manufacturer’s protocol.

### Electroporation of cells

Electroporations were carried out as described by Thibault *et al*. (27). Both Huh7.5 miR-122 KO cells and Huh 7.5 Drosha KO cells were electroporated using the following conditions: 225 V, 950 μF, 4 mM, and ∞ Ω.

### Transient HCV replication assay

On Day 0 Huh-7.5 cells were co-electroporated as described previously with 5ug of J6/JFH-1 Rluc wild-type RNA or mutant viral RNAs and 60 pmol of miR-122 or control microRNA into approximately 6.0 × 10^6^ cells in 400 μL Dulbecco’s phosphate-buffered saline (PBS). Electroporated cells were resuspended in 4 ml cell culture medium and 500 μL of cells were plated in 6 well dishes and incubated at 37°C to be harvested at different time points. The electroporation protocol for Huh 7.5 Drosha KO cells is similar to described above, however, a total of 750 uL of resuspended electroporated cells are plated on the 6 well dishes instead of 500 uL.

### Transient HCV replication assay with preknockdown

Exonuclease (XRN1) and Pyrophosphatases (DOM3Z, DUSP11) preknockdown assay were performed as described previously (28). 48 hours before day 0, approximately 6.0 × 10^6^ cells in 400 μL Dulbecco’s PBS were electroporated with 60 pmol of each siRNA together for preknockdown and were plated in 15-cm tissue culture dishes, two cuvettes per dish. siRNA-treated cells were incubated for 2 days at 37°C. On day 0, cells were collected from the dishes, prepared as described for electroporation (two cuvettes’ worth from each dish), and then electroporated with viral RNA, siRNAs, and microRNAs and plated as described above.

### Luciferase assay

To measure luciferase expression cells were washed with Dulbecco’s phosphate-buffered saline and the lysate was harvested by using 100 μL of passive lysis buffer. Renilla or dual-luciferase kits (Promega, Madison, USA) were used to measure luciferase as suggested by the manufacturer’s protocols. 10 μL of lysate was mixed with the appropriate luciferase assay substrate and light emission was measured by using a Glomax 20/20 Luminometer (Promega, Madison, USA).

### Translation assay

10 μg of J6/JFH-1 Rluc2a GNN viral RNA (WT/mutants) and 1ug of control T7 Fluc mRNA were co electroporated in Huh 7.5 miR-122 KO cells. Cells were harvested at 2 hours post electroporation and a dual-luciferase test was done to analyze Rluc and Fluc expression.

### Secondary structure prediction

5’ UTR 1-117 nucleotide of the positive strand and 3’UTR 1-117 of the negative strand of the HCV genome were used for RNA structure prediction. RNA prediction software ‘RNA structure’ from Matthews lab at https://rna.urmc.rochester.edu/index.html was used for RNA structure prediction (52). Dot-bracket files were generated using Fold and bifold algorithms for predicting the structure of single RNA or two interacting RNAs respectively. VARNA (VARNA GUI applet) was used to generate the folded RNA images from the dot-bracket files (53).

### Data analysis

All experiment results are displayed as averages from at least three independent experiments. Error bars indicating standard errors of the mean (SEM). Statistical analyses were performed using GraphPad Prism 7.4 (San Diego USA); statistical significance was determined by the tests indicated in each figure legend.

## FUNDING

This work was supported by grants from the Canadian Institutes of Health Research [MOP-133458], the Canada Foundation for Innovation [18622], and the University of Saskatchewan (CoMBRIDGE) to J.A.W. M.P. was funded by a University of Saskatchewan Graduate Teaching Fellowship, and M.A.P was funded by a Natural Sciences and Engineering Research Council of Canada, MSc Scholarship, a Canadian Network on Hepatitis C (CanHepC) Training Program, Doctoral Research Scholarship, and by a University of Saskatchewan CoMGRAD Scholarship.

## ACKNOWLEDGMENTS

We would like to acknowledge Charlie Rice (The Rockefeller University) for providing the J6/JFH-1(p7Rluc2A) and Huh-7.5, and Drosha KO Huh 7.5 cells, and Matthew Evans (Icahn School of Medicine at Mount Sinai) for Huh-7.5 miR-122 KO cells.

## AUTHOR CONTRIBUTIONS

M.P, M.A.P, and J.A.W, designed and performed the experiments, and analyzed the data; M.P and J.A.W. wrote the manuscript.

---

**FIG S1**: Analysis of a mutant virus with an insertion into SLII of J6/JFH-1 (p7Rluc2a) (Ins Mutant)

Structure prediction and Translation assay of Ins Mutant RNA compared with wild-type HCV in the presence or absence of miR-122, and 265-67G-C is a translation defective mutant used as a negative control. The lowest free energy structure prediction models of Ins mutant with or without miR-122 are presented. The structures were predicted using the ‘RNA structure’ online tool and the first favorable lowest free energy structure is presented (52). These structures are prediction models and have not been experimentally validated. Error bar indicates the standard deviation of the mean and asterisk indicates significant differences. The significance was determined by using one-way ANOVA (*P<0.033, **P<0.002, ***P<0.001). Predicted structure of Ins mutant with and without miR-122.

**FIG S2**: Structure prediction of 1-117 nucleotide of J6/JFH-1 (p7Rluc2a) wild-type HCV and mutants with all possible mutations at position 25.

Wild-type HCV with U at 25th position predicts to fold to the canonical SLII structure when miR-122 is annealed to the sequence. Similar to wild-type HCV with miR-122, mutant HCV with C, A, and G at the 25^th^ position is also predicted to form the canonical SLII structure. Wild-type HCV without miR-122 is predicted to form a thermodynamically stable SLII^Alt^ structure. The structures were predicted using the ‘RNA structure’ online tool and the first favorable lowest free energy structure is presented. These structures are prediction models and have not been experimentally validated.

**FIG S3**: Structure prediction models of J6/JFH-1 (p7Rluc2a) miR-122 binding site 1 and site 2 HCV mutants.

1-117 nucleotide sequence of these mutants were analysed using the ‘RNA structure’ online tool, and the 4 lowest free energy structure models are presented. These structures are prediction models and have not been experimentally validated.

**FIG S4**: RNA structure prediction models of J6/JFH-1 (p7Rluc2a) miR-122 binding site 2 HCV mutants (HCV-S2-GGCGUG and HCV-S2C-GUGAGG).

(A) 1-117 nucleotide sequences from the 5’ UTR of the genomic RNA are analyzed using ‘RNA structure’ online tool, and the 4 lowest free energy structure models are presented. (B) 1-105 nucleotide sequence from the complementary sequence on the 3’ UTR of the negative strand RNA of the wild-type virus, HCV-S2-GGCGUG, HCV-S2C-GUGAGG, and S2p5 (C39G) are analyzed using the same online tool, and the most favorable lowest free energy structure models are presented. These structures are prediction models and have not been experimentally validated.

